# Combating the SARS-CoV-2 Omicron variant with non-Omicron neutralizing antibodies

**DOI:** 10.1101/2022.01.30.478305

**Authors:** Yingdan Wang, Xiang Zhang, Jiangyan Liu, Yanqun Wang, Wuqiang Zhan, Mei Liu, Meng Zhang, Qimin Wang, Qianying Liu, Tongyu Zhu, Yumei Wen, Zhenguo Chen, Jincun Zhao, Fan Wu, Lei Sun, Jinghe Huang

**Affiliations:** Key Laboratory of Medical Molecular Virology (MOE/NHC/CAMS) and Shanghai Institute of Infectious Disease and Biosecurity, the Fifth People’s Hospital of Shanghai, Shanghai Public Health Clinical Center, Institutes of Biomedical Sciences, School of Basic Medical Sciences, Fudan University, Shanghai 200032, China; State Key Laboratory of Respiratory Disease, National Clinical Research Center for Respiratory Disease, Guangzhou Institute of Respiratory Health, the First Affiliated Hospital of Guangzhou Medical University, Guangzhou, Guangdong 510182, China; Institute of Infectious Disease, Guangzhou Eighth People’s Hospital of Guangzhou Medical University, Guangzhou, Guangdong 510060, China

**Keywords:** COVID-19, SARS-CoV-2, Neutralizing Ab, Omicron Variant

## Abstract

The highly mutated and transmissible Omicron variant has provoked serious concerns over its decreased sensitivity to the current coronavirus disease 2019 (COVID-19) vaccines and evasion from most anti-severe acute respiratory syndrome coronavirus 2 (SARS-CoV-2) neutralizing antibodies (NAbs). In this study, we explored the possibility of combatting the Omicron variant by constructing bispecific antibodies based on non-Omicron NAbs. We engineered ten IgG-like bispecific antibodies with non-Omicron NAbs named GW01, 16L9, 4L12, and REGN10987 by fusing the single-chain variable fragments (scFvs) of two antibodies through a linker and then connecting them to the Fc region of IgG1. Surprisingly, eight out of ten bispecific antibodies showed high binding affinity to the Omicron receptor-binding domain (RBD) and exhibited extreme breadth and potency against pseudotyped SARS-CoV-2 variants of concern (VOCs) including Omicron, as well as authentic Omicron(+R346K) variants. Six bispecific antibodies containing the cross-NAb GW01 neutralized Omicron variant and retained their abilities to neutralize other sarbecoviruses. Bispecific antibodies inhibited Omicron infection by binding to the ACE2 binding site. A cryo-electron microscopy (cryo-EM) structure study of the representative bispecific antibody FD01 in complex with the Omicron spike (S) revealed 5 distinct trimers and one unique bi-trimer conformation. The structure and mapping analyses of 34 Omicron S variant single mutants elucidated that two scFvs of the bispecific antibody synergistically induced the RBD-down conformation into 3-RBD-up conformation, enlarged the interface area, accommodated the S371L mutation, improved the affinity between a single IgG and the Omicron RBD, and hindered ACE2 binding by forming bi-trimer conformation. Our study offers an important foundation for anti-Omicron NAb design. Engineering bispecific antibodies based on non-Omicron NAbs may provide an efficient solution to combat the Omicron variant.

## INTRODUCTION

Omicron, first identified in South Africa and reported to the WHO at the end of Nov. 2021, became the dominant severe acute respiratory syndrome coronavirus 2 (SARS-CoV-2) variant globally in Jan. 2022. Omicron is the most severely altered version of SARS-CoV-2, with more than 30 mutations in the spike protein, sixteen of which are in the receptor-binding domain (RBD). Omicron variants extensively escape neutralization by sera from vaccinated or convalescent individuals ^1–10^. Moreover, most neutralizing antibodies (NAbs), including many clinical-stage monoclonal antibodies (mAbs), have completely lost their neutralization potency against Omicron ^1,3,4,11^. Therefore, there is an urgent need to explore and develop countermeasures against the Omicron variant.

In this study, we used four NAbs named GW01, 16L9, 4L12, and REGN10987, which failed to bind or neutralize the Omicron variant, to engineered full-length IgG-like bispecific antibodies. We surprisingly found that these bispecific antibodies could neutralize all the VOCs, including Omicron and authentic Omicron(+R346K), while the parental antibody cocktail showed no neutralization against Omicron. Cryo-EM structure study showed six dynamic states of the Omicron S trimer upon bispecific antibody binding, including a novel bi-trimer conformation, within which RBDs were all in “up” conformations. This bi-trimer is critical for inhibiting ACE2 binding and explains the superiority of the bispecific antibody. These novel bispecific antibodies are strong candidates for the treatment and prevention of infection with the Omicron variant and VOCs and other sarbecoviruses that may cause future emerging or reemerging coronavirus diseases.

## RESULTS

### Isolation of three non-Omicron neutralizing antibodies from COVID-19 convalescent individuals

We sorted and cultured SARS-CoV-2 S-specific memory B cells from two recovered coronavirus disease 2019 (COVID-19) patients and discovered three anti-SARS-CoV-2 NAbs, designated GW01, 4L12, and 16L9. The germlines and CDR3 of these antibodies are listed in Table S1. All three antibodies showed strong binding to the RBD of SARS-CoV-2 (**Fig. 1A**). However, they had no or weak binding to the S trimer or S RBD of the Omicron variant (**Fig. 1A**).

**Figure 1.**
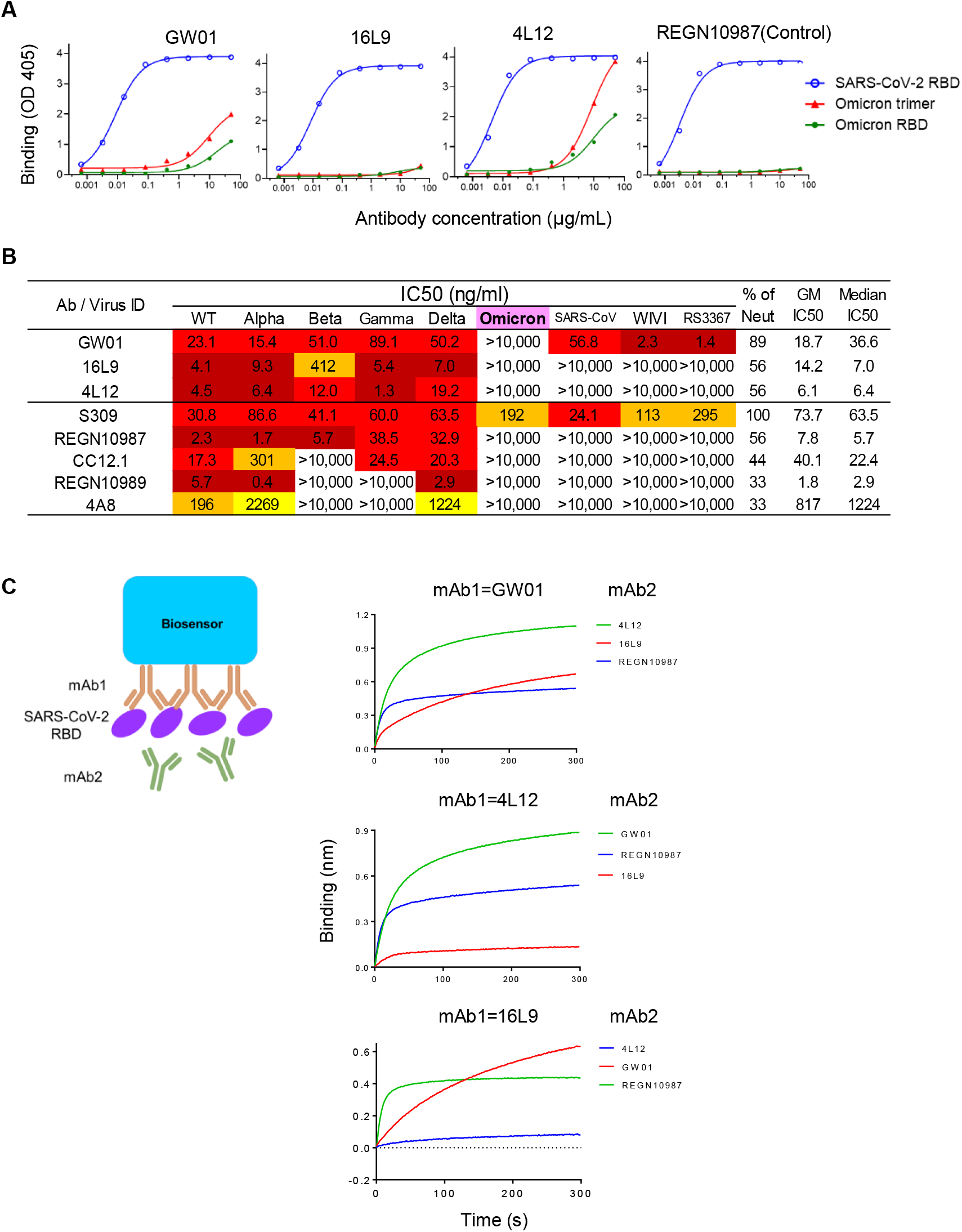
Isolation of three non-Omicron neutralizing antibodies from COVID-19 convalescent individuals. **(A)** Binding of GW01, 16L9, and 4L12 to the SARS-CoV-2 RBD, Omicron RBD and trimer in an ELISA. REGN10987 was used as a control. **(B)** Neutralizing activities of GW01, 16L9, and 4L12 were determined against pseudotyped SARS-CoV-2 and its variants Alpha, Beta, Gamma, Delta, and Omicron, as well as sarbecoviruses. REGN10987 was used as a control. **(C)** Binding of 4L12, 16L9, and RGN10987 to the SARS-CoV-2 RBD in competition with GW01, as measured by bilayer interferometry experiments.

GW01, 4L12, and 16L9 potently neutralized SARS-CoV-2 and the VOCs Alpha, Beta, Gamma, and Delta, but they failed to neutralize the Omicron variant (**Fig. 1B**). A panel of control NAbs failed to neutralize the Omicron except S309. S309 neutralized Omicron to a similar degree as previous reports^12^,^13^. GW01 was a cross-NAb that was able to neutralize SARS-CoV and the SARS-related coronaviruses (SARSr-CoVs) RS3367 and WIV1. GW01 antibodies showed no competition with 4L12, 16L9 or the control antibody REGN10987 in binding the RBD (**Fig. 1C**), indicating that GW01 binds to an epitope different from that bound by 4L12, 16L9, and REGN10987.

### Binding and neutralization of the Omicron variant by bispecific antibodies

We constructed bispecific antibodies targeting different epitopes in the RBD using GW01 in combination with 16L9, 4L12, and REGN10987, and explored their possibilities to neutralize Omicron variant. We linked the single-chain variable fragments (scFvs) of the parental antibodies with a (Gly4Ser)4 linker and then fused them to a hinge-CH2-CH3 fragment of human immunoglobulin (hIgG1 Fc) to generate a single gene-encoded IgG-like bispecific antibody (**Fig. 2A**). For example, the sequence order of the GW01-16L9 (FD01) bispecific antibody was as follows: GW01 VL-(Gly_4_Ser)_3_-GW01 VH-(Gly4Ser)4-16L9 VL-(Gly_4_Ser)_3_-16L9 VH-hinge-CH2-CH3. SDS–PAGE results showed that the size of the single chain of two representative bispecific antibodies, GW01-16L9 and 16L9-GW01, was approximately 100 kDa and that the purity was >95% (**Fig. 2B**). Crosslinking bispecific antibodies with glutaraldehyde revealed that the full size of the bispecific antibodies was approximately 200 kDa, which was 10% larger than that of the parental antibodies (180 kDa, **Fig. 2B**).

**Figure 2.**
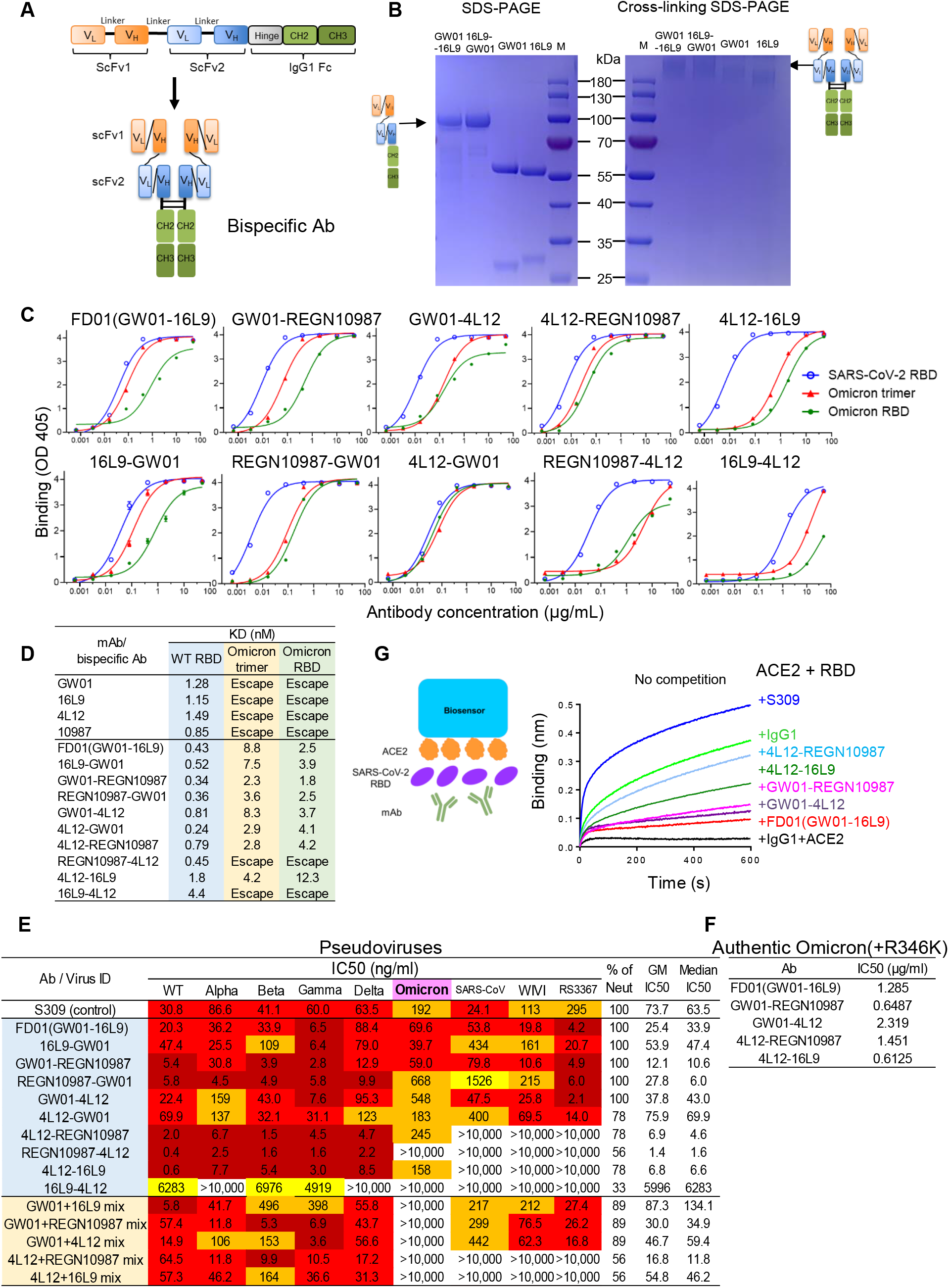
Binding and neutralization of the Omicron variant by bispecific antibodies. **(A)** Schematic diagrams showing the structures of bispecific antibodies. **(B)** SDS–PAGE and cross-linking SDS–PAGE gels showing the sizes of the representative bispecific antibodies and their parental antibodies. **(C)** Binding specificities of the bispecific antibodies to the SARS-CoV-2 RBD-his, Omicron trimer-his, or RBD-his protein. **(D)** Binding affinities of the bispecific antibodies to the SARS-CoV-2 RBD-his, Omicron trimer-his, or RBD-his protein were measured by bilayer interferometry experiments. **(E)** Neutralization by bispecific antibodies and combinations of parental antibodies against the VOCs, including Omicron variant, and sarbecoviruses. **(F)** Neutralization of five representative bispecific antibodies against authentic Omicron(+R346K) variants. **(G)** Five representative bispecific antibodies bind to the ACE2-binding site and block RBD binding to ACE2. Binding of ACE2 to the SARS-CoV-2 RBD in competition with bispecific antibodies (red), S309 (blue), control IgG1 (green), and IgG1+ACE2 (black).

We constructed ten bispecific antibodies and tested their binding abilities to the RBD or S trimer of SARS-CoV-2 and the Omicron variant. Eight bispecific antibodies, FD01 (GW01-16L9), 16L9-GW01, GW01-REGN10987, REGN10987-GW01, GW01-4L12, 4L12-GW01, 4L12-REGN10987, and 4L12-16L9, not only strongly bound to the RBD of SARS-CoV-2 and the S trimer and RBD proteins of the Omicron variant (**Fig. 2C**) but also showed high binding affinity to these proteins (**Fig. 2D**). These results indicated that the structure of the bispecific antibody increased the parental antibody binding affinity to the Omicron RBD.

To understand the breadth of these bispecific antibodies, we performed a neutralization assay using SARS-CoV-2 pseudoviruses, including Alpha, Beta, Gamma, Delta, and Omicron variants, and the sarbecoviruses SARS-CoV, WIV1 and RS3367. Surprisingly, these eight bispecific antibodies potently neutralized the Omicron variant with IC50 values from 39.7 to 548 ng/ml. Six bispecific antibodies containing the cross-NAb GW01 strongly neutralized all the tested VOCs and sarbecoviruses (**Fig. 2E**). FD01 and GW01-REGN10987 were the best broadly NAbs, with geometric mean (GM) IC50 values of 25.4 and 12.1 ng/ml, respectively. 4L12-REGN10987 and 4L12-16L9 strongly neutralized all the tested VOCs with GM IC50 values of 6.9 and 6.8 ng/ml, respectively (**Fig. 2E**). However, the parental NAb combinations showed no neutralization against the Omicron variant (**Fig. 2E**). Taken together, these data indicated that bispecific antibodies consisting of non-Omicron NAbs efficiently neutralize the Omicron variant in a way that is different from the antibody cocktail.

To confirm the neutralization efficacy of the bispecific antibodies, we performed plaque reduction neutralization assays with an authentic Omicron variant containing the R346K mutation, which escapes more SARS-CoV-2 NAbs than the Omicron variant^3^. All five representative bispecific antibodies efficiently neutralized the live Omicron variant (**Fig. 2F**), confirming that the bispecific antibodies composed of non-Omicron NAbs are able to neutralize the Omicron variant.

### Bispecific antibodies bind to the ACE2-binding site

Competition assays were then performed to evaluate the abilities of the bispecific antibodies to inhibit the binding of SARS-CoV-2 RBD to the recombinant ACE2 protein (**Fig. 2G**). All five representative bispecific antibodies prevented RBD binding to ACE2 protein, while the control antibody S309 did not affect the S/ACE2 interaction. FD01 showed a strong inhibitory effect against ACE2 binding. These results indicated that bispecific antibodies inhibit Omicron variant infection by occupying the ACE2-binding site on the RBD.

### Snapshots of Omicron S-FD01 structures determined by cryo-EM

To further investigate the neutralization mechanism of the bispecific antibodies, we chose FD01 (GW01-16L9) as a representative antibody for structural study. Local refinement focused on the RBD and ScFvs improved the interface region to 3.51 Å resolution and allowed us to unambiguously build the RBD and scFvs (**Table S2**). We determined the cryo-EM structure of the prefusion stabilized SARS-CoV-2 Omicron S ectodomain trimer in complex with the bispecific antibody FD01 (Omicron S-FD01), revealing 6 states of the complex: In the state 1 Omicron S-FD01 structure (~26% of the particles), only one 16L9 binds to the “up” RBD, and the two down RBDs have no antibody binding (up-down-down RBDs, 1 scFv, 3.47 Å). In state 2 (~38%), the bispecific antibody FD01 (GW01-16L9) binds to a widely open RBD, the so-called “wide_up” state, 16L9 binds to an “up” RBD, and the third RBD remains in a down state (wide_up-up-down RBDs, 3 scFvs, 3.70 Å). In both state 3 (~8%) and state 4 (~15%), two FD01 (GW01-16L9) bind separately to two “wide_up” state RBDs, and the third RBD represents the half-up conformation in state 3 (wide_up-wide_up-half_up RBDs, 4 scFvs, 3.91 Å) and the up conformation in state 4 (wide_up-wide_up-up RBDs, 4 scFvs, 3.47 Å), without antibody binding. In state 5 (~4%), all RBDs are in the “wide_up” state, each bound with an FD01 (GW01-16L9) (all wide_up RBDs, 6 scFvs, 3.87 Å). In the final state 6 (~4% of the particles), two state 5 trimers are connected by three bispecific FD01 antibodies, forming a bi-trimer structure (bi-trimer, all wide_up RBDs, 12 scFvs, 6.11 Å) (**Fig. 3, Fig. S2 and S3**).

**Figure 3.**
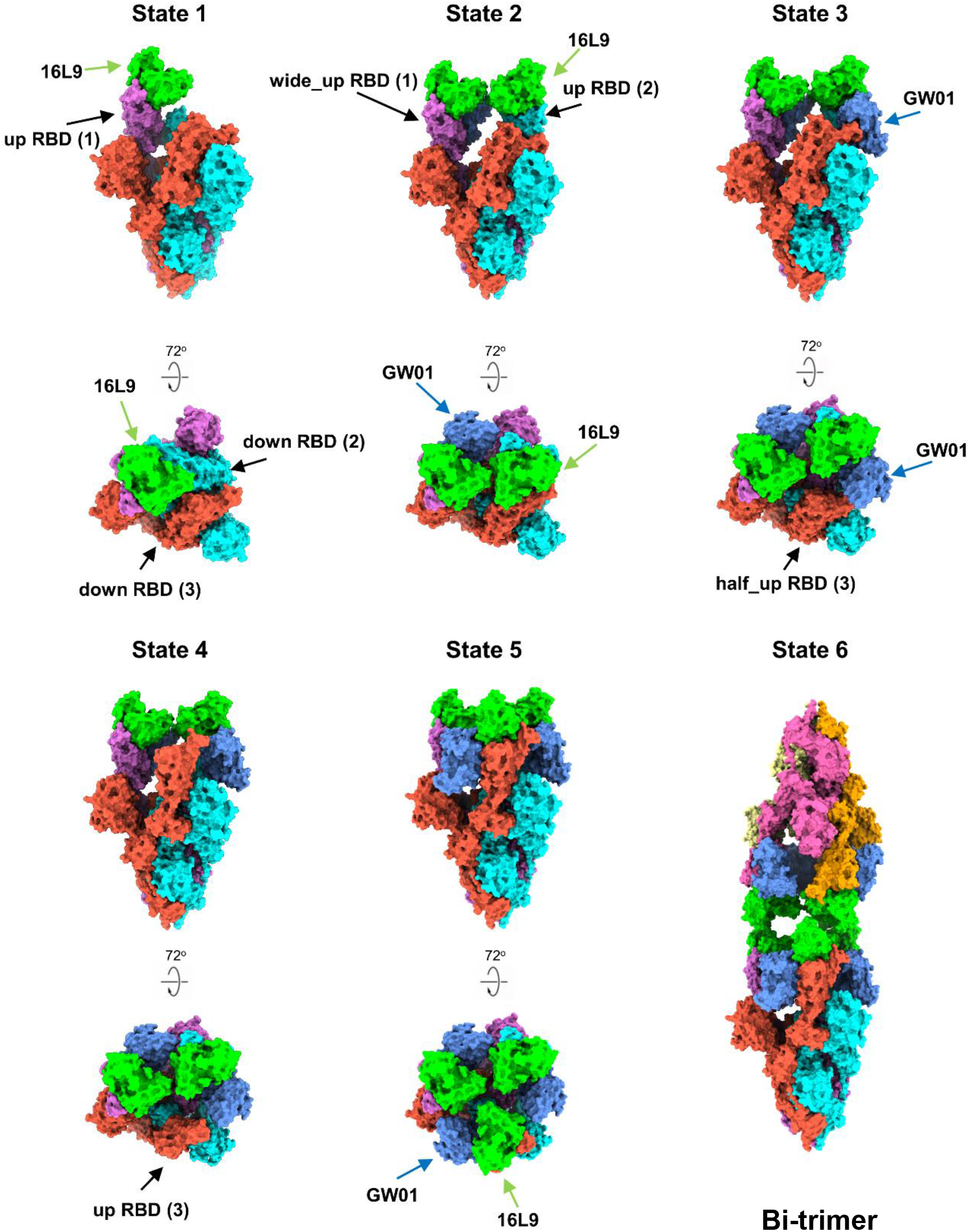
Cryo-EM structures of the Omicron S trimer in complex with the bispecific antibody FD01. The bispecific antibody FD01 binds to Omicron S trimers in six states. Two perpendicular views of Omicron S-FD01 are shown in surface representation, with 16L9 ScFv in lime and GW01 ScFv in cornflower blue.

### Collaborative binding mechanism of FD01 bispecific antibody

These six cryo-EM structures represent the conformational transitions of the Omicron S trimer upon FD01 binding. To simplify the presentation, three protomers of a spike trimer are clockwisely defined as 1, 2, and 3 (**Fig. 3, Fig. 4**). The apo spike trimer (state 0) includes one regular up RBD and two down RBDs. First, 16L9 binds to the “up” RBD (RBD-1) of the apo spike trimers and forms the state 1 confirmation. After that, GW01, connected with 16L9, binds to RBD-1, inducing it into a wide_up state (via an ~13 Å outward motion) and pushing RBD-2 to flip from the “down” to the “up” state, making enough space to accommodate the first bispecific antibody FD01 (16L9 and GW01) on RBD-1 and the 16L9 of the second FD01 on RBD-2. A slight ~3 Å inward motion of RBD-3 is induced by the neighboring RBD-2 to stabilize this state (state 2). Then, in state 3, the “up” RBD-2 opens up further to the “wide_up” state, allowing both 16L9 and GW01 of the second FD01to bind RBD-2. In addition, RBD-3 is pushed up to a “half_up” state. In state 4, the “half_up” RBD-3 opens up to the regular “up” state and is ready for 16L9 binding. Following that, in state 5, the third FD01 binds to RBD-3 and induces RBD-3 to adopt the “wide_up” state. Finally, two trimers in state 5 form a bi-trimer induced by three pairs of Fc regions from six antibodies.

**Figure 4.**
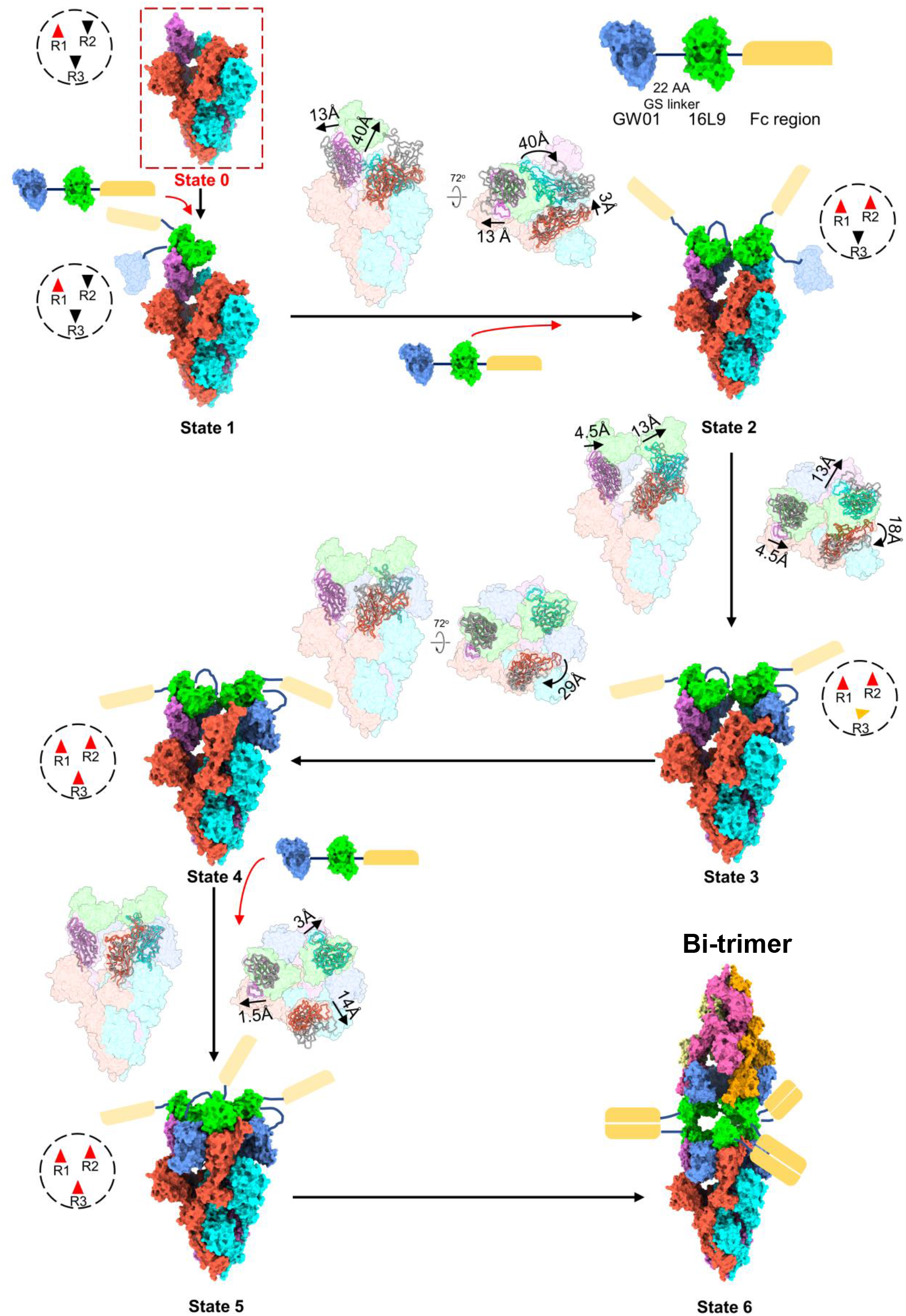
Conformation transitions of Omicron S-FD01 in all states. State 0 inside the red dashed box is a hypothetical apo state structure. Small triangles inside the black dashed circle indicate the up/half_up/down states of three RBDs in the trimer.

Thus, the six states represent the continuous conformational transitions starting from the first 16L9 binding to three FD01 binding and the final bi-trimer formation, which inhibits the ACE2 binding by aggregating virions. In addition to the motion of RBDs, the N-terminal domains (NTDs) of the trimer are also moved or rotated following the motion of their neighboring RBDs.

### FD01 targets two conserved epitopes

16L9 and GW01 bind to two different sites of one RBD (**Fig. 5A**). The epitope of 16L9 almost overlaps with the receptor-binding motif (RBM), while GW01 binds outside the RBM. The binding of 16L9 and the RBD buries a 1061 Å^2^ surface area, and a total of 20 residues from RBD are involved. The interaction between 16L9 and RBD is largely driven by extensive hydrophilic and hydrophobic interactions between CDRH1, CDRH2, CDRH3 and CDRL1 of 16L9 and RBD (**Fig. 5C**). Residues D405, T415, D420, Y421, Y453, L455, F456, Y473, A475, N477, Y489, R493, P500, Y501, and H505 of the RBD are involved in this interaction, forming 14 pairs of hydrogen bonds and 3 patches of hydrophobic interactions (**Fig. 5C**). In addition, the hydrogen bond between S96 of CDRL3 and R403 from the RBD and the salt bridge between E52 of CDRL2 and R493 from the RBD further enhance the interaction (**Fig. 5C**). Coincidentally, residues Y453, A475, Y489, R493, T500, Y501, and H505 of the Omicron RBD are important for ACE2 recognition and binding^14^. The neutralization activity test showed that 16L9 alone was able to broadly neutralize SARS-CoV-2 and SARS-CoV-2 variants. This result implies that 16L9 targets the conserved residues of the RBM, which is needed for receptor binding.

**Figure 5.**
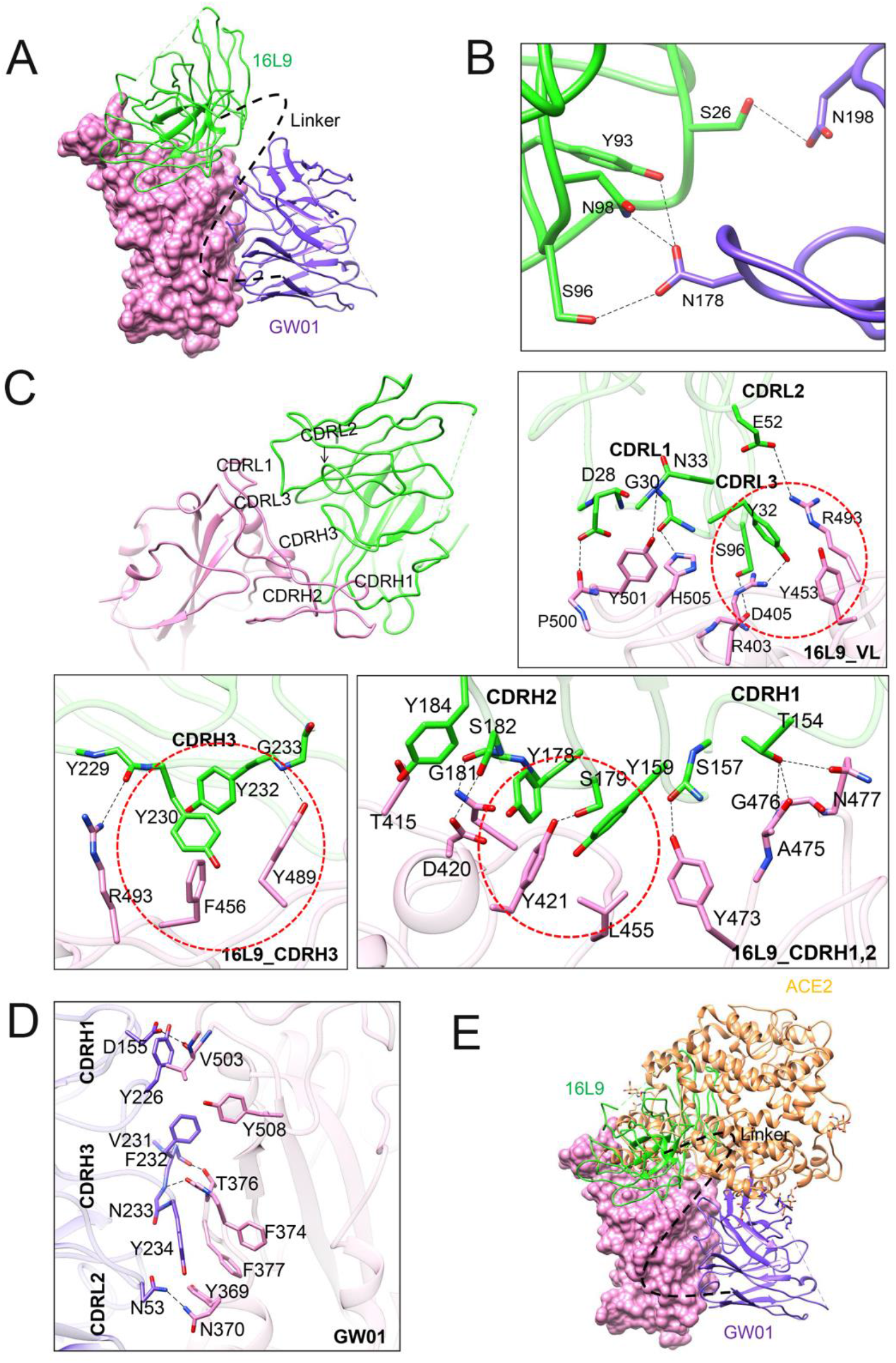
Two conserved epitopes recognized by FD01. (**A**) Close-up view of the interaction between FD01 and Omicron; the Omicron RBD is displayed in pink in surface representation. 16L9 and GW01 are shown as cartoons colored green and medium-blue, respectively. (**B**) The interface between 16L9 and GW01. (**C-D**) The interaction of 16L9 (**C**) and GW01 (**D**). The residues involved in interactions are represented as sticks. Polar interactions are indicated as dotted lines. (**E**) Ribbon diagrams of FD01 and ACE2 (PDBID: 7T9L) bound to the Omicron RBD.

GW01 interacts with another novel conserved epitope beyond the binding site of 16L9. The binding site of GW01 and the RBD has a buried surface area of 668.2 Å. The interaction between GW01 and the RBD is mainly contributed by CDRH3. The long loop (226-YGPPDVFNY-234) of CDRH3 engages with Y369, F374, T376, F377, Y508, and V503 from the RBD, forming 3 patches of hydrophobic interactions and 2 pairs of hydrogen bonds. D155 on CDRH1 and N53 on CDRL2 are also involved in the interaction by forming hydrogen bonds between V503 and N370, respectively (**Fig. 5D**).

Interestingly, the simultaneous binding of 16L9 and GW01 with the RBD introduces additional interactions. Hydrogen bonds are formed between N178 and N196 of GW01 and S26, Y93, S96 and N98 of 16L9 (**Fig. 5B**), which further enhances the interaction between FD01 and S.

Structural alignment of the 16L9-GW01-RBD complex with the ACE2–RBD complex indicated that both 16L9 and GW01 were able to compete with ACE2 when binding to the RBD (**Fig. 5E**), which is consistent to the competition assay.

### Bispecific antibodies accommodated the mutations in the Omicron variant

We constructed 34 single mutants of the Omicron variant to identify the key residues that mediate resistance to GW01, 16L9, 4L12, and REGN0987. The S371L mutation, which was found to stabilize the Omicron into a single-RBD-down conformation^12^, greatly decreased the neutralization activities of GW01, 4L12, and REGN10987 (**Table 1**). The S375F mutation decreased the neutralization activity of GW01 by 16-fold and resulted in resistance to REGN10987. The K417N mutation resulted in complete resistance to 16L9 (>1000-fold). All six tested bispecific antibodies showed only a slight decrease (12.4-to 25.5-fold) or no change in neutralization activity against the S371L mutant. Therefore, bispecific antibodies bind to the ACE2-binding site of the RBD and accommodate the S371L mutation of the Omicron variant, resulting in extraordinary breadth for these bispecific antibodies.

**Table 1.**
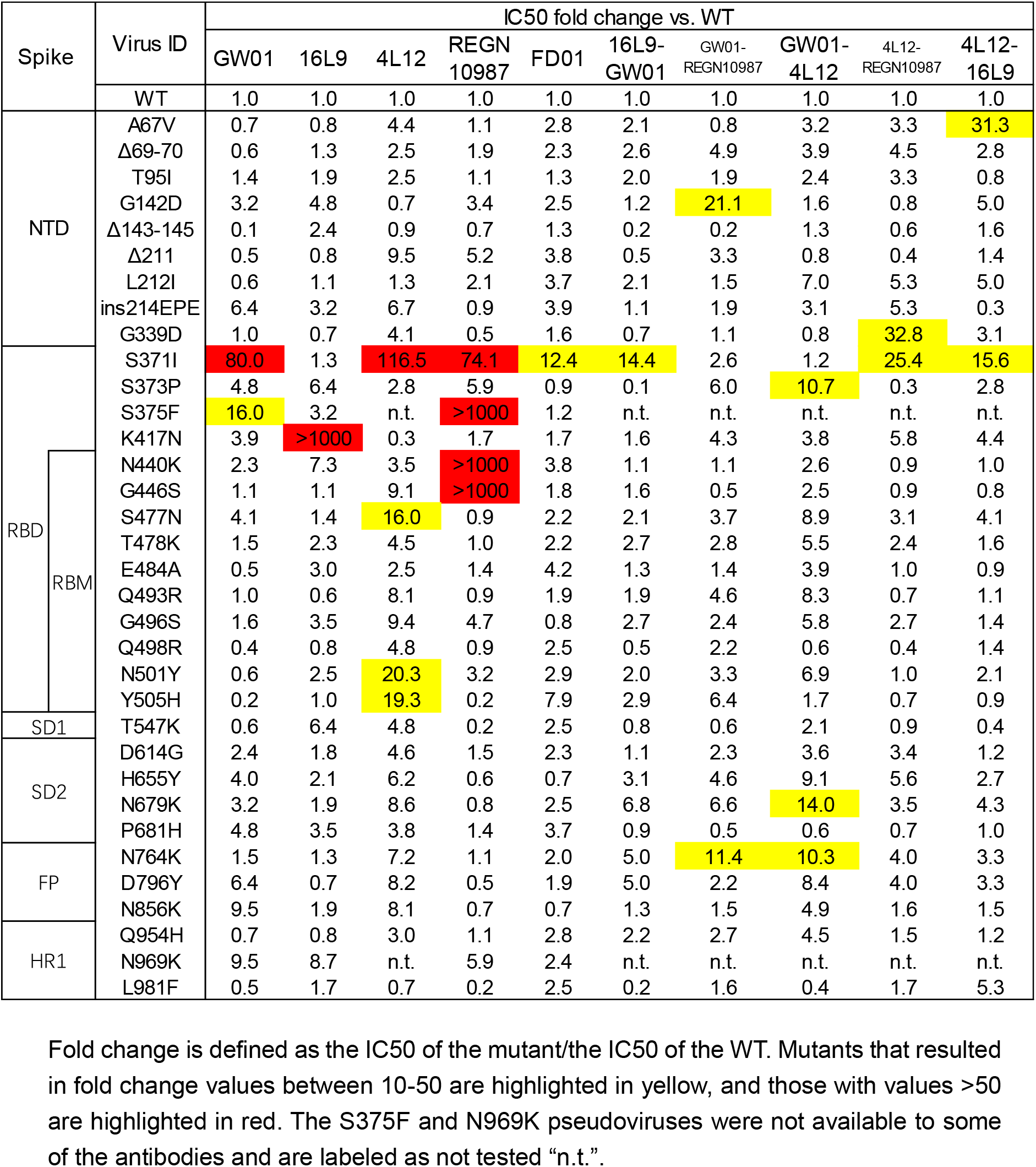
Neutralization by GW01, 16L9, 4L12, REGN10987, and six bispecific antibodies against 34 strains containing single mutations present within the Omicron variant.

## DISCUSSION

The recently emerged SARS-CoV-2 Omicron strain raised unprecedented global concern about invalidation of most FDA-approved antibody drugs, including LY-CoV555, LY-CoV016, REGN10933, REGN10987, AZD8895 and AZD1061^15^. Previous studies have shown that combining two NAbs that target different neutralizing epitopes in the SARS-CoV-2 S protein increased therapeutic and prophylactic efficacy. However, the most potent anti-SARS-CoV-2 antibody cocktail, RENG10933/REGN10987, failed to neutralize the Omicron variant. Zhou tested 10 antibody combinations against Omicron and three antibody combinations with increasing neutralization contained a VH1-58 derived anti-Omicron NAb that bound RBD in the “up” position. Cyro-EM structure showed that the antibody combination with improved neutralization (B1-182.1/A19-46.1) synergistically induced the 3-RBD-up conformation^12^. However, this antibody combination approach is not an ideal solution because few anti-Omicron NAbs are available.

Using the NAbs that failed to neutralize the Omicron variant, we constructed a serial of novel bispecific antibodies that capable of neutralizing all SARA-CoV-2 variants of concern (VOCs), including the Omicron strain. Interestingly, single IgG parental antibodies or the combination of parental antibodies failed to neutralize Omicron, although the neutralization activity against SARS-CoV-2 or SARS-CoV-2 Alpha, Beta, Gamma, and Delta was remarkable. Thus, the effective neutralization of Omicron by bispecific antibodies may be due to the construction of this bispecific antibody.

The structure of FD01 (GW01-16L9) bispecific antibody gives a hit at this collaborative binding mechanism. First, one 16L9 scFv binds to the exposed epitope of the “up” state RBD-1 in the apo Omicron S trimer as a trigger. Then, the 20 aa GS linker between 16L9 and GW01 guides GW01 to its targeting epitope of RBD-1, pushing RBD-1 more open, which unlocks the “down” state of RBD-2 and induces it to adopt the up state. Thus, another 16L9 scFv could easily catch the “up” state RBD-2. The same triggering process would occur on RBD-2 and RBD-3, allowing the binding of the second and the third FD01 to RBD-2 and RBD-3, respectively. GW01 Fab alone could not trigger binding to the apo state of the Omicron trimer without the help of 16L9 and the GS linker guider. Although 16L9 can bind to the first “up” state RBD, it lacks the ability to release other “down” state RBDs for further binding (**Fig. S5**). In summary, the two scFvs of the GW01-16L9 bispecific antibody have collaborative roles in the neutralization process. The neutralization mechanism of FD01 may be mediated by the unique engineering of the combination of two antibodies into one, which induces RBD-up conformation, enlarges the interface area, improves the affinity of a single IgG and the RBD, stabilizes the interaction by additional interactions between the two antibodies, forms bi-trimer which hinders the ACE2 binding by aggregating virions, and therefore blocks the infection of SARS-CoV-2 Omicron variant.

Taken together, the unique construction of bispecific antibodies enables non-Omicron NAbs to neutralize Omicron variant. Our approach can rescue the majority of the SARS-CoV-2 antibodies, such as REGN10987, to overcome the resistance of Omicron and prepare for future SARS-CoV-2 variants.

## ACKNOWLEDGMENTS

We thank Center of Cryo-Electron Microscopy, Fudan University for the supports on cryo-EM data collection. This work was supported by the National Natural Science Foundation of China (31771008 to JH and 81900729 to LS), the National Major Science and Technology Projects of China (2017ZX10202102 to JH and 2018ZX10301403 to FW), Shanghai Municipal Health Commission (2018BR08 to JH), the Chinese Academy of Medical Sciences (2019PT350002 to JH).

## AUTHOR CONTRIBUTIONS

JH, LS, and FW conceived and designed the experiments. JH, ML, and FW performed B cell sorting and antibody cloning. YDW and YJ constructed the bispecific antibodies and performed neutralization assay, ELISA, bilayer interferometry experiments. YDW, YJ, and QW constructed and expressed SARS-CoV-2 pseudovirus mutants and purification antibodies. LS, XZ, WZ, and ZC performed the structural studies. JZ and YQW were responsible for the authentic virus experiments. JH, YDW, YJ, FW, LS, XZ, WZ, JZ, and YQW analyzed the data. YW and TZ supervised the project. JH, LS, FW, YDW, and XZ wrote the manuscript.

## COMPETING INTERESTS

Patents about the bispecific antibodies in this study are pending.

## Materials and Methods

### Cell lines, proteins, viruses and plasmids

The human primary embryonic kidney cell lines (HEK293T) and 293T-hACE2 cells were cultured in DMEM medium with 10% fetal bovine serum (FBS). RBD-his proteins of SARS-CoV-2 and Omicron variant were purchased from Sino Biological. Genes of bispecific antibodies were synthesized by Genscript. The authentic Omicron (B.1.1.529) with R346K mutation used in this study were isolated from COVID-19 patients in Guangzhou, passaged, and titered on Vero E6 cells. African green monkey kidney-derived Vero E6 cells were grown in Dulbecco’s modified Eagle’s medium (DMEM; Gibco, Grand Island, NY, USA) supplemented with 10% fetal bovine serum (FBS). All work with authentic SARS-CoV-2 was conducted at the Guangzhou Customs Technology Center Biosafety Level 3 (BSL-3) Laboratory.

### Production of Pseudoviruses

S genes of SARS-CoV-2 (NC_045512), Alpha (containing 69–70 and 144 deletions and N501Y, A570D, D614G, P681H, T716I, S982A, and D1118H substitutions), Beta (containing D80A, D215G, 241-243 deletions and K417N, E484K, N501Y, D614G and A701V substitutions), Gamma (containing L18F,T20N,P26S,D138Y, R190S,K417T, E484K,N501Y, D614G.H655Y,T1027I, and V1176F substitutions), Delta (containing T19R, 157-158 deletions and L452R, T478K, D614G, P681R and D950N substitutions), and Omicron (containing A67V, 69-70del, T95I, G142D, 143-145del, N211I, 212del, ins215EPE, G339D, S371L, S373P, S375F, K417N, N440K, G446S, S477N, T478K, E484A, Q493R, G496S, Q498R, N501Y, Y505H, T547K, D614G, H655Y, N679K, P681H, N764K, D796Y, N856K, Q954H, N969K, L981F substitutions), SARS-CoV, bat SARSr-CoVs (WIV1 and Rs3367) were synthesized by BGI and constructed in pcDNA3.1 vector. Pseudoviruses were generated by co-transfection of 293T cells with an env-deficient HIV backbone pNL4-3.Luc.R-E-backbone and a spike expressing vector^16^,^17^. Fifty additional spike variants carrying currently circulating single-point mutations were constructed by site-directed mutagenesis.

### Neutralization assay

The neutralization activities of mAbs and bispecific antibodies were determined using a single-round pseudovirus infection of 293T-hACE2 cells. 10 μl of 5-fold serially diluted antibody was incubated with 40 μl of pseudovirus in 96-well plate at 37 °C for 1 h. 10^4^ 293T-hACE2 cells were then added to the mixture and cultured for 48 h at 37 °C. Cells were lysed and firefly luciferase activity were developed with a luciferase assay system (Promega) and detected on a luminometer (Perkin Elmer). The IC50s of NAbs were calculated using the GraphPad Prism 7.04 software (La Jolla, CA, USA).

### Memory B-cell staining, sorting and antibody cloning

CD19+IgA–IgD–IgM-primary B cells were sorted out from peripheral blood mononuclear cells (PBMC) of recovered patients of COVID-19 and expanded *in vitro* in MEM medium with 10% FBS in the presence of irradiated 3T3-msCD40L feeder cells, IL-2 and IL-21 as previously described ^18^. After fifteen days of incubation, supernatants were screened for neutralization against SARS-CoV-2. From the wells with SARS-CoV-2 neutralization activities, the variable regions of the antibody (VH and VL) genes were amplified by RT–PCR. mAbs were expressed as human IgG1 by HEK293F cells and purified using a protein G column (Smart-Lifesciences).

### ELISA

2 μg/ml of SARS-CoV-2 RBD-his, Omicron trimer-his, and Omicron RBD-his protein were coated overnight at 4 °C in a 96-well plate (MaxiSorp Nunc-immuno, Thermo Scientific, USA). Wells were blocked with 5% non-fat milk (Biofroxx, Germany) in PBS for 1 hour at room temperature, followed by incubation with 5-fold serially diluted mAb in disruption buffer (PBS, 5% FBS, 2% BSA, and 1% Tween-20) for 1 hour at room temperature. After 3 washing steps with PBS 0.05% Tween 20 (PBS-T), 1:2500 diluted HRP-conjugated goat anti-human IgG antibody (Jackson Immuno Research Laboratories, USA) was added for 1 hour at room temperature. Plates were washed three times with 0.2% Tween-20 in PBS and developed using ABST (Thermo Scientific, USA) for 30 minutes. Absorbance at 405 nm was read on a Multiskan FC plate reader (Thermo Scientific, USA).

### Biolayer interferometry (BLI) binding assay and competition assay

Experiments were carried out on a FortéBio OctetRED96 instrument. The kinetics of monoclonal antibody binding to SARS-CoV-2 RBD-his, Omicron RBD-his, or trimer-his proteins was measured using anti-human IgG (AHC) biosensors. 10 μg/ml of mAbs were immobilized on biosensors for 200s. After a 120 sec stabilization step with 0.02% PBST (PBS with 0.02% Tween), biosensors were moved into 6 μg/ml of RBD-his or trimer-his proteins for the 300 sec association step. Then biosensors were moved into 0.02% PBST to detect dissociation for 300 sec. The buffer control binding was subtracted to deduct nonspecific binding. K_on_, K_off_, and KD were calculated by FortéBio Data Analysis software (Version 8.1) using 1:1 binding and a global fitting model.

For the Ab competition assay, 10 μg/ml of mAbs 1 were immobilized on the anti-human IgG (AHC) biosensors for 200s. After wash with 0.02% PBST for 120s to reach baseline, biosensors were moved into 50 μg/ml of IgG1 isotype control for 200s and then moved into SARS-CoV-2 RBD at 6 μg/ml for 300s. After wash with 0.02% PBST for 120s, biosensors were moved into 10 μg/ml of mAb2 for 600s to detect the association between mAb2 and SARS-CoV-2 RBD.

For ACE2 competition, biosensors were moved into 20 μg/ml of ACE2-Fc for 600s. After baseline, wash, and blocking steps, biosensors were moved into pre-mix of 600 nM of mAb and 100 nM SARS-CoV-2 RBD for 600s. A mixture of ACE2-Fc and SARS-CoV-2 RBD was used as a positive control, while the mixture of IgG1 isotype control and SARS-CoV-2 RBD was used as a negative control.

### Focus reduction neutralization test

SARS-CoV-2 Omicron(+R346K) focus reduction neutralization test (FRNT) was performed in a certified Biosafety level 3 lab. Fifty microliters antibody were serially diluted, mixed with 50 μl of SARS-CoV-2 (100 focus forming unit, FFU) in 96-well microwell plates and incubated for 1 hour at 37°C. Mixtures were then transferred to 96-well plates seeded with Vero E6 cells (ATCC, Manassas, VA) for 1 hour at 37°C to allow virus entry. Inoculums were then removed before adding the overlay media (100 μl MEM containing 1.2% Carboxymethylcellulose, CMC). After 24-hour post infection, the overlay was discarded and the cell monolayer was fixed with 4% paraformaldehyde solution for 2 hours at RT. After permeabilized with 0.2% Triton X-100 for 20 min at room temperature, the plates were sequentially stained with cross-reactive rabbit anti-SARS-CoV-2 N IgG (Cat. No.: 40143-T62, Sino Biological Inc) as the primary antibody and HRP-conjugated goat anti-rabbit IgG(H+L) (No.: 109-035-088, Jackson ImmunoResearch) as the secondary antibody in 37°C for 1 hour respectively. The reactions were developed with KPL TrueBlue Peroxidase substrates. The numbers of SARS-CoV-2 foci were calculated using CTL ImmunoSpot S6 Ultra reader (Cellular Technology Ltd). Neutralizing activity was defined as the ratio of inhibition of SARS-CoV-2 focus comparing diluted antibody to control.

### Construction and expression of bispecific NAbs

Genes of a bispecific Ab consisting of the scFv of GW01 and scFv of 16L9, REGN10987 or 4L12 were synthesized and codon-optimized by GenScript. The bispecific antibody sequence alignment was as follows: variable light chain (VL) and variable heavy chain (VH) of mAb 1 or mAb 2 were linked with a (Gly_4_Ser)_3_ linker. VL-VH of mAb 1 and VL-VH of mAb 2 were linked with a (Gly4Ser)4 linker and then fused to the expression vector with hinge-CH2-CH3 fragment of human immunoglobulin (hIgG1 Fc). FD01 bispecific antibody sequence order was as follows: GW01 VL-(Gly_4_Ser)_3_-GW01 VH-(Gly4Ser)4-16L9 VL-(Gly_4_Ser)_3_-16L9 VH-hinge-CH2-CH3.

293F cells were transiently transfected with bispecific Abs plasmid. After 6 days of culture at 37°C in a 5% CO2 incubator, supernatant was collected and filtered. Bispecific antibodies were purified with protein G colume (Smart-Lifesciences) and stored in PBS at −80°C.

### SDS-PAGE and cross-linking SDS-PAGE of bispecific antibodies

The purity and molecular weight of bispecific antibodies were then analyzed by SDS-PAGE and cross-linking SDS-PAGE. Briefly, 5 μg of bispecific antibodies were mixed with 5x SDS-loading sample buffer containing 10% β-mercaptoethanol. The samples were heated for 10 min at 100°C and were then loaded on an SDS gradient gel (4–20% Precast Protein Improve Gels, Genscript Biotech Corporation). The gel was run for 120 min at 120 V, and Coomassie staining was performed.

Extent of dimer was investigated by cross-linking of bispecific antibodies with glutaraldehyde (Sigma-Aldrich). Briefly, 5 μg of antibodies were diluted in 25 μl of PBS in the presence of a 2.7 μM of glutaraldehyde cross-linker. The mixture was incubated at RT for 5 minutes, and then glutaraldehyde was quenched by adding 1 M Tris-HCl buffer (pH 8.0) to a final concentration of 40 mM. After mixing with 5x SDS-loading sample, the protein samples were loaded on a 4-20% SDS gradient gel. The gel was run for 180 min at 120 V and confirmed by Coomassie staining.

### Expression and purification of SARS-CoV-2 Omicron Spike

The Human codon gene encoding SARS-CoV-2 Omicron S ectodomain was purchased from GeneScript. The expression plasmid of Omicron S 6P substitution^19^ was constructed and transfected into suspension HEK293F using polyethlenimine. After 72 hours, the supernatants were harvested and filtered for affinity purification by Histrap HP (GE). The protein was then further purified by gel filtration using Superose 6 increase 10/300 column (GE Healthcare) in 20 mM Tris pH 8.0, 200 mM NaCl.

### Cryo-EM sample preparation

Purified SARS-CoV-2 Omicron S at 1.554 mg/mL was mixed with FD01 antibody by a molar ratio of 1:1.5 incubated for 10 min on ice before application onto a freshly glow-discharged holey amorphous nickel-titanium alloy film supported by 400 mesh gold grids ^20^. The sample was plunged freezing in liquid ethane using Vitrobot IV (FEI/Thermo Fisher Scientific), with 2 s blot time and −3 blot force and 10 s wait time.

### Cryo-EM data collection and image processing

Cryo-EM data were collected on a Titan Krios microscope (Thermo Fisher) operated at 300 kV, equipped with a K3 summit direct detector (Gatan) and a GIF quantum energy filter (Gatan) setting to a slit width of 20 eV. Automated data acquisition was carried out with SerialEM software^21^ through beam-image shift method^22^.

Movies were taken in the super-resolution mode at a nominal magnification 81,000×, corresponding to a physical pixel size of 1.064 Å, and a defocus range from −1.2 μm to −2.5 μm. Each movie stack was dose-fractionated to 40 frames with a total exposure dose of about 58 e^-^/Å^2^ and exposure time of 3s.

All the data processing was carried out using either modules on, or through, RELION v3.0^23^ and cryoSPARC^24^. A total of 4,363 movie stacks was binned 2 × 2, dose weighted, and motion corrected using MotionCor2 ^25^ within RELION. Parameters of contrast transfer function (CTF) were estimated by using Gctf ^26^. All micrographs then were manually selected for further particle picking upon ice condition, defocus range and estimated resolution.

Remaining 3,817 good images were imported into cryoSPARC for further patched CTF-estimating, blob-picking and 2D classification. From 2D classification, bi-trimer and trimer particles were observed. Several good 2D classes of these two kind particles were used as templates for template-picking separately. After 2D classification of particles from template-picking was finished, all good particles from blob-picking and template-picking were merged and deduplicated, subsequently being exported back to RELION through pyem package ^27^.

For bi-trimer map, 1,003,956 particles were extracted at a box-size of 540 and rescaled to 180, then carried on 2 round of 3D classification with a soft circular mask of 480 Å in diameter in RELION. Only good classes were selected, yielding 166,441 clean particles. These particles were re-extracted unbinned (1.064 Å/pixel) and auto-refined without applying symmetry, yielding a map at 6.11 Å.

For trimer map, 1,003,956 particles were extracted at a box-size of 320 and rescaled to 160, then carried on 1 round of 3D classification with a soft circular mask of 220 Å in diameter in RELION. Three classes with different conformation change on trimer RBDs were selected separately for another round of 3D classification. Particles in different states were auto-refined, CTF-refined and polished separately. Some density of RBDs and Fabs in some state were not well-resolved, so we carried out no-alignment 3D classification with NTD-RBD-Fabs (NRF) mask to improve those regions. Finally, we got 5 states of Omicron S-FD01 trimer.

To get clear interfaces of RBD with Fabs, we did local-refinement focused on that region. We first selected all good 3D-classes with relatively complete RBD and FD01 density within all states. We auto-refined these particles with a C3-aligned reference, but the auto-refinement procedure was not applied any symmetry. Then particles were expanded with C3 symmetry and further subtracted with one NRF mask. After no-alignment 3D-classification, 249,122 particles with complete NRF density were selected out, exported to cryoSPARC and carried out local refinement, yielding a local-refined map at 3.51 Å.

The reported resolutions above are based on the gold-standard Fourier shell correlation (FSC) 0.143 criterion. All the visualization and evaluation of 3D density maps were performed with UCSF Chimera ^28^ and ChimeraX^29^. The above procedures of data processing are summarized in Fig. S3 and Fig. S4. These sharpened maps were generated by DeepEMhancer ^30^ and then “vop zflip” to get the correct handedness in UCSF Chimera for subsequent model building and analysis.

### Model building and refinement

For model building of SARS-CoV-2 Omicron S FD01 complex, the SARS-CoV-2 Omicron S trimer model and the antibody model generated by swiss-model ^31^ were fitted into the map using UCSF Chimera and then manually adjusted with COOT ^32^. Several iterative rounds of real-space refinement were further carried out in PHENIX ^33^. The RBD bounded with 16L9 and GW01 was refined against the local refinement map and then docked back into global refinement trimer and bi-trimer maps. Model validation was performed using phenix.MolProbity. Figures were prepared using UCSF Chimera and UCSF ChimeraX^29^.

### Data and materials availability

The cryo-EM map and the coordinates of SARS-CoV-2 Omicron S complexed with FD01 have been deposited to the Electron Microscopy Data Bank (EMDB) and Protein Data Bank (PDB) with accession numbers EMD-32655 and PDB 7WOQ (state 1), EMD-32656 and PDB 7WOR (state 2), EMD-32657 and PDB 7WOS (state 3), EMD-32659 and PDB 7WOU (state 4), EMD-32660 and PDB 7WOV (state 5), EMD-32661 and PDB 7WOW (state 6), EMD-32654 and PDB 7WOP (NTD-RBD-GW01-16L9 local refinement).

---

**Table S1.** The germline and CDRH3 sequences of GW01, 4L12, and 16L9.

| Donor ID | mAb ID | VH | CDRH3 sequence | VL | CDRL3 sequence |
| --- | --- | --- | --- | --- | --- |
| Donor 1 | GW01 | IGHV3-43 | AKDRSYGPPDVFNYEYGMDV | IGLV1-44 | AAWDDSLNWV |
| Donor 1 | 4L12 | IGHV3-66 | ARDLITYGMDV | IGKV1-9 | QQLNSYPPLT |
| Donor 2 | 16L9 | IGHV3-53 | ARGEIQPYYYYGMDV | IGLV2-8 | SSYAGSSNFDV |

**Table S2.**
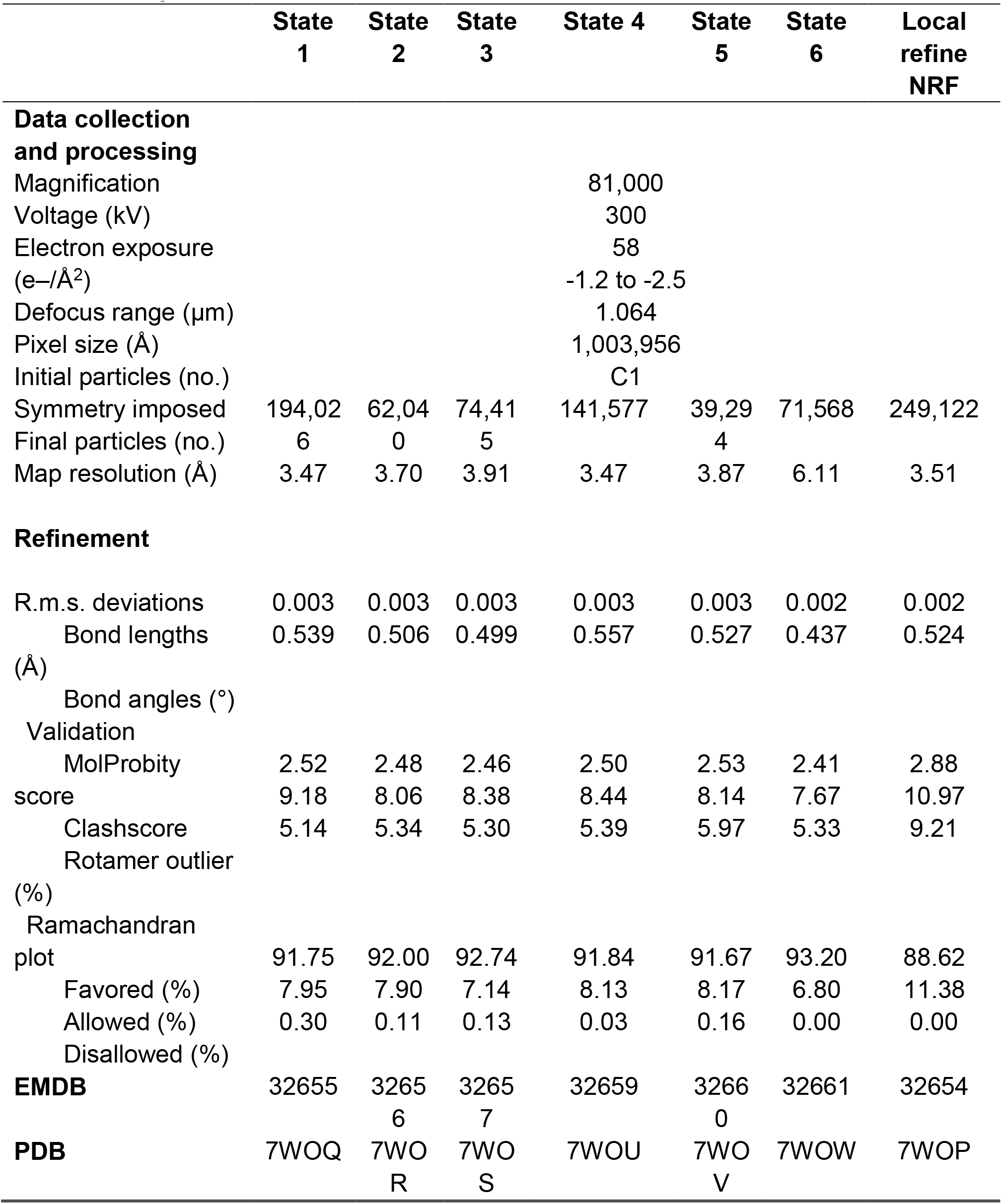
Cryo-EM data collection and refinement statistics.

**Figure S1.**
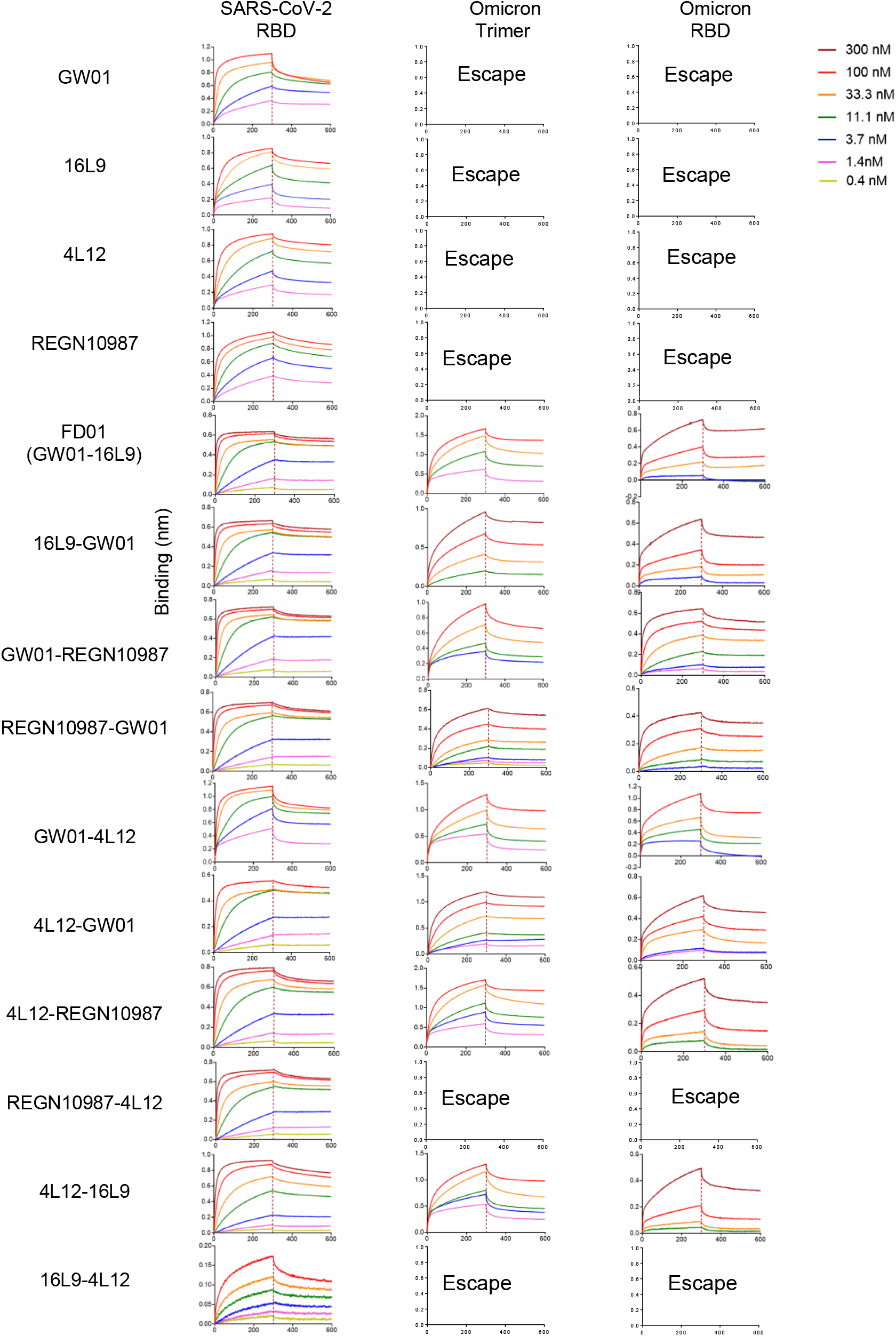
Binding affinities of GW01, 16L9, 4L12, REGN10987, and ten bispecific antibodies to SARS-CoV-2 RBD-his, Omicron trimer-his and Omicron RBD-his measured by bilayer interferometry experiments. Antibodies were immobilized on anti-human IgG (AHC) biosensors and then tested for their binding abilities to the target proteins.

**Figure S2.**
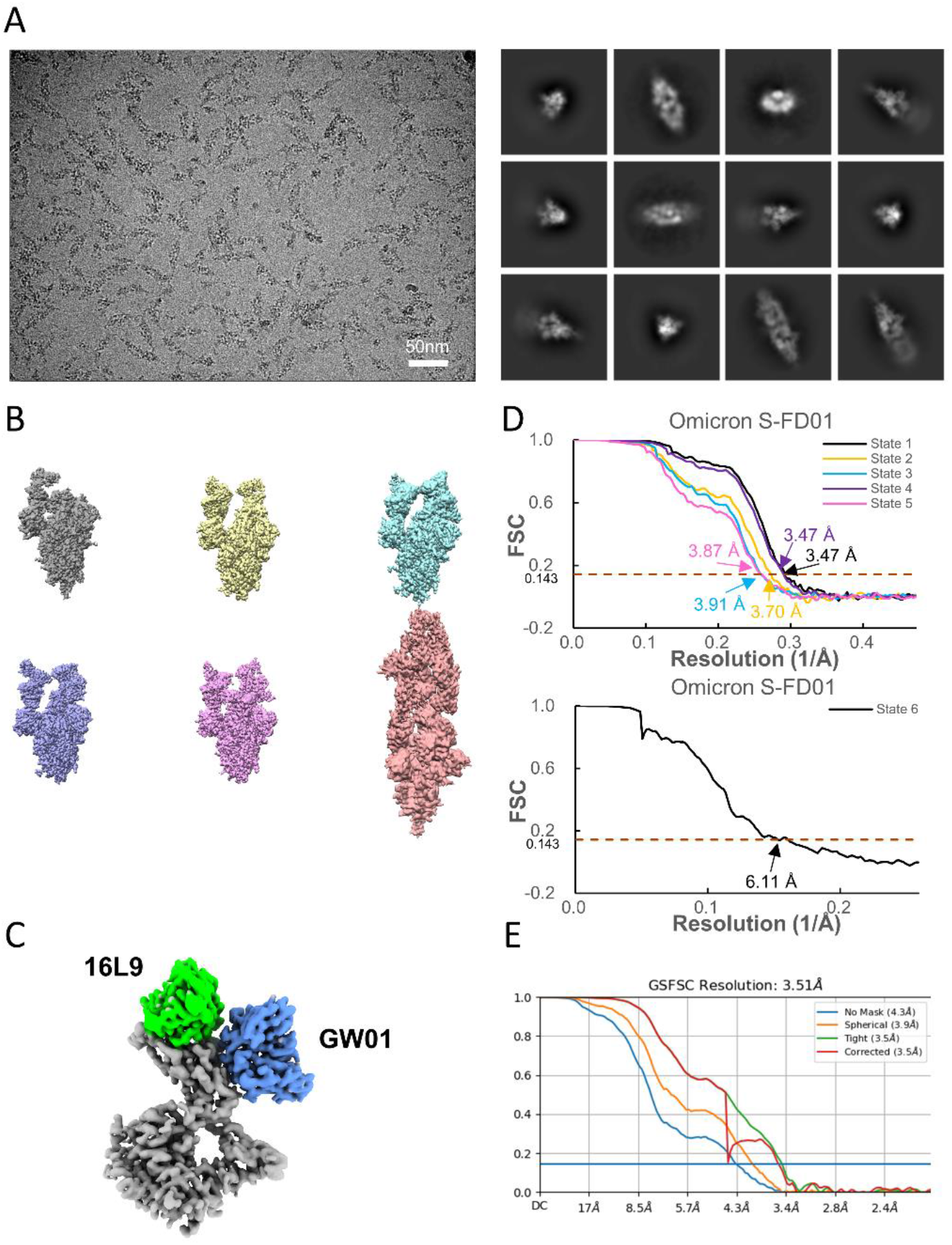
Cryo-EM data collection and processing of FD01 bound SARS-CoV-2 Omicron S. **(A)** Representative electron micrograph and 2D classification results of FD01 bound SARS-CoV-2 S. **(B)** The reconstruction map of the complex structures at six states. **(C)** The local-refined map of the NRF region. **(D)** Gold-standard Fourier shell correlation curves generated in RELION for structures of six states. The 0.143 cut-off is indicated by a horizontal dashed line. **(E)** Gold-standard Fourier shell correlation curves generated in cryoSPARC for local-refined map.

**Figure S3.**
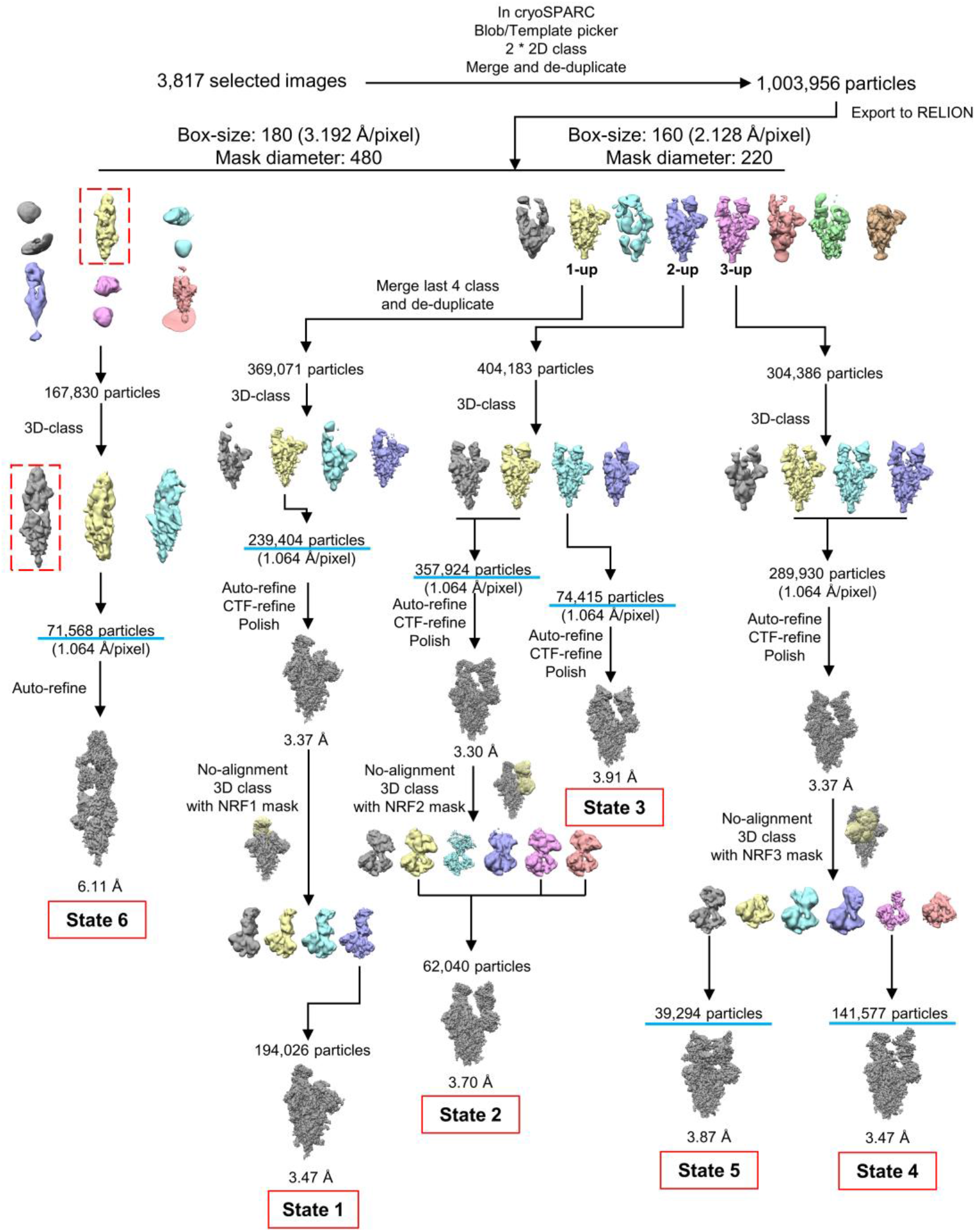
Data processing flowchart of FD01 bound SARS-CoV-2 Omicron S trimer. Particles number above cyan line is used for particle counting statistics.

**Figure S4.**
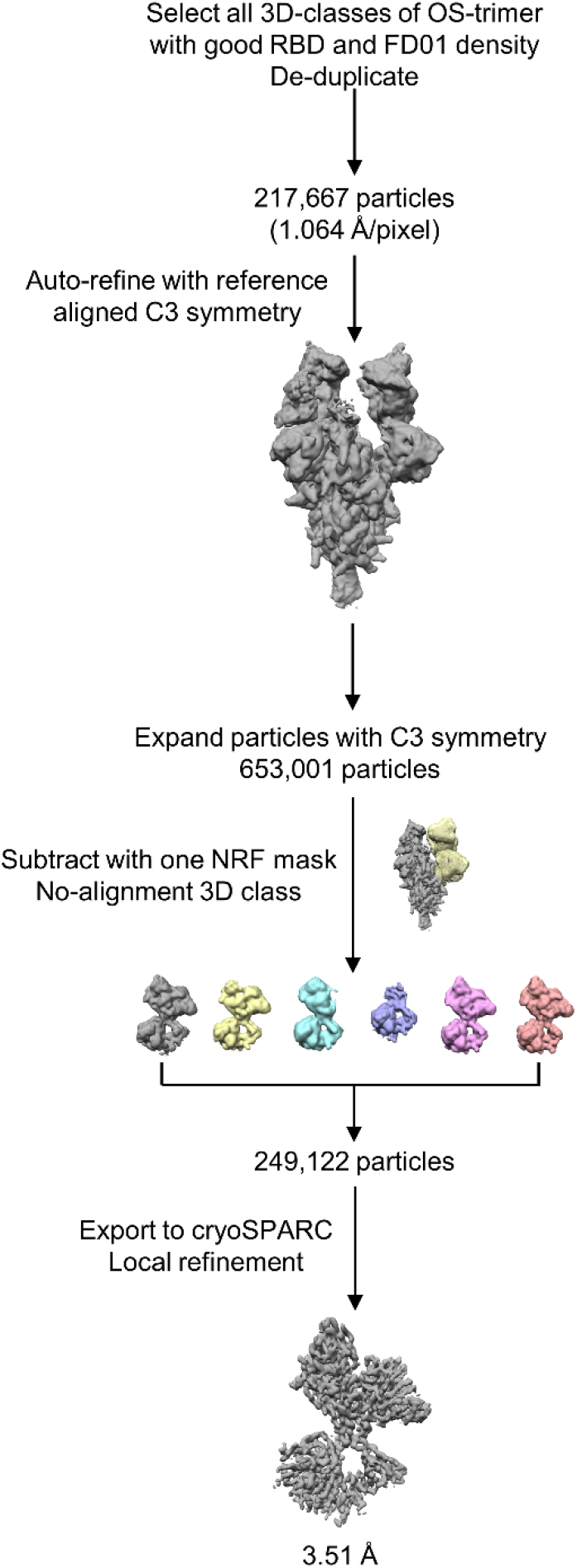
Data processing flowchart of local refinement of RBD-FD01.

**Figure S5.**
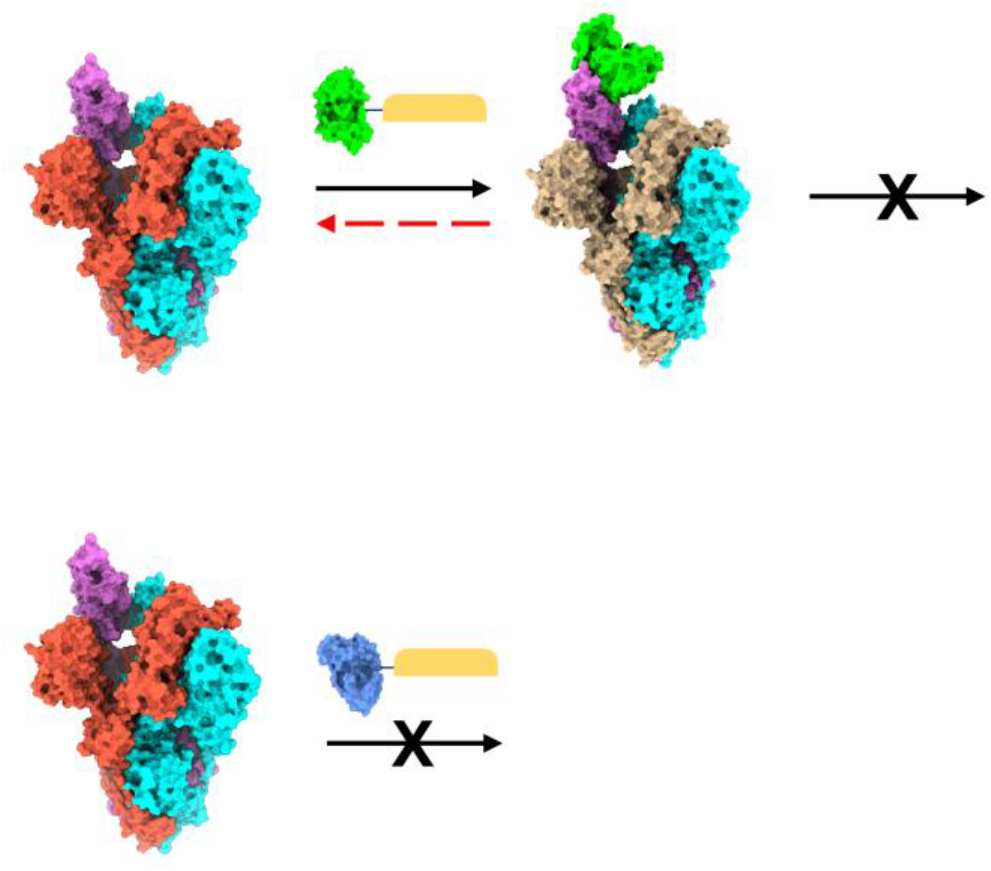
Hypothesis of binding features when Omicron S trimer meets with mAbs of 16L9 or GW01.

## Notes

### Competing Interest Statement

The authors have declared no competing interest.

